# Loss of intracerebellar heterogeneity and selective vulnerability in Spinocerebellar ataxia type 1 neurodegeneration

**DOI:** 10.1101/2022.02.24.481789

**Authors:** Katherine Hamel, Emmanuel Labrada Moncada, Carrie Sheeler, Juao-Guilherme Rosa, Stephen Gilliat, Ying Zhang, Marija Cvetanovic

**Author notes:** Corresponding Author: Marija Cvetanovic, Department of Neuroscience, University of Minnesota, 2101 6^th^ Street SE, Minneapolis, MN 55455.

## Abstract

Regional heterogeneity of neurons and glia is a key feature of the brain, yet the effect of disease on heterogeneity and its relationship with selective neuronal vulnerability remains poorly understood.

Using region-specific RNA sequencing, we identified a large number of differentially expressed genes (DEGs) across distinct regions of the cerebellar cortex, supporting the notable intrinsic regional transcriptional heterogeneity of the healthy cerebellum. Further, we used fiber photometry to identify regional physiological differences in the activity of Purkinje cells (PCs) during self-motivated, unrestrained walking and non-walking states. In the inherited cerebellar neurodegenerative disease Spinocerebellar ataxia type 1 (SCA1), patients exhibit preferential degeneration of the posterior cerebellum, suggesting regionally selective vulnerability. We demonstrated that in a mouse model of SCA1 the Purkinje cells and glia residing in the posterior vermis of cerebellum also undergo earlier and more severe pathology. Intriguingly, the intrinsic transcriptional heterogeneity of anterior and posterior cerebellum seen in healthy mice was diminished in SCA1 mice. This disruption was also demonstrated via fiber photometry, where we found notable impacts in PC activity in the posterior cerebellum as well as loss of regional differences in PC activity during self-motivated, unrestrained walking, and non-walking states in SCA1 mice. Our findings indicate regionally distinct mechanisms of pathogenesis across cerebellar regions that result in reduced intracerebellar heterogeneity.

## Introduction

Heterogeneity of cell types throughout and within brain regions is a hallmark of a healthy brain. Selective vulnerability, or severe loss of neurons in a disease-specific brain region, characterizes all neurodegenerative diseases (Fu et al., 2018; Pandya and Patani, 2021; Saxena and Caroni, 2011) (Subramaniam, 2019)(Fu et al., 2018). The role of heterogeneity in normal brain function and in region specific vulnerability in neurodegenerative diseases, and how disease impacts heterogeneity are areas of particular interest to the neuroscience community and key questions of modern neuroscience (Pandya and Patani, 2021).

The cerebellum contains the majority of the brain’s neurons. Although the cerebellumis largely known for its role in movement and balance, its role in non-motor behavior such as cognition, affect, and reward is increasingly recognized in the field. Anatomically, the cerebellar cortex is divided into two lateral hemispheres and a central vermis, which is distinguished by the primary fissure into an anterior and posterior area (D’Mello and Stoodley, 2015). While cerebellar circuitry looks almost identical across these regions, several studies have demonstrated differences in gene expression, function, and connectivity across these cerebellar regions (Buckner et al., 2011; Choe et al., 2018; Kozareva et al., 2021; Middleton and Strick, 1994; Shakkottai et al., 2017; Zhou et al., 2014).

Spinocerebellar ataxia type-1 (SCA1) is an inherited neurodegenerative disease that severely affects the cerebellum and for which there is no disease modifying therapy currently available. SCA1 belongs to a group of polyglutamine (polyQ) diseases that also includes Huntington’s disease, SCA, 2, 3, 6, 7, and 17, Spinal and Bulbar Muscular Atrophy (SBMA), and Dentatorubral-Pallidoluysian Atrophy (DRPLA) (Orr et al., 1993). In SCA1, expansions of 39 or more CAG repeats in the *ATAXIN1* (*ATXN1)* gene result in expression of a mutant polyglutamine (polyQ) ATXN1 protein (Zoghbi and Orr, 2009). While mutant ATXN1 is expressed throughout the brain, SCA1 is characterized by a severe loss of cerebellar Purkinje cells (PCs) (Klinke et al., 2010; Koeppen, 2005; Seidel et al., 2012; Zoghbi et al., 1988). Moreover, pathological studies have revealed that the greatest PC loss occurs in the posterior cerebellar vermis of patients with SCA1 (Reetz et al., 2013). A recent longitudinal MRI study indicated significant pathology in the posterior vermis, in particular in lobule X, and in hemispheres, with relative sparing of anterior lobules of SCA1 patients (Nigri et al., 2022).This suggests two possibilities-either that the PCs of the posterior vermis and hemispheres are selectively vulnerable to ATXN1-driven pathology and/or that PCs of the anterior vermis are uniquely resistant to ATXN1 pathology. Investigating these mechanisms will increase our understanding of pathogenesis and may inspire development of new SCA1 therapies.

Here, we comprehensively explored intrinsic cerebellar heterogeneity at the level of gene expression and PC activity during unrestricted movement in healthy mice. To increase our understanding of heterogeneity and selective vulnerability in cerebellar disease, we used the *Atxn1^154Q/2Q^* mice, in which 154 CAG repeats were knocked-in into one endogenous mouse *Atxn1* allele (Watase et al., 2002). *Atxn1^154Q/2Q^* mice have been extensively used to study SCA1 pathogenesis, but most of the pathological studies have been focused on the primary fissure of the vermis (lobules IV/V and VI) and/or on bulk cerebellar RNA or protein analysis (Duvick et al., 2010; Ingram et al., 2016; Ruegsegger et al., 2016). Using immunohistochemistry, we extensively examined SCA1 neuronal and glial pathology throughout the cerebellum. We then used region-specific bulk RNAsequencing to explore the molecular underpinnings of intracerebellar heterogeneity and selective vulnerability across the lateral hemispheres and anterior and posterior vermis in SCA1 mice. Finally, we determined how SCA1 alters the intrinsic activity of anterior and posterior PCs in freely moving mice using fiber photometry.

## Materials and Methods

### Mice

The creation of the *Atxn1^154Q/2Q^* mice was previously described (Watase et al., 2002). These knock-in mice express mutant protein under the endogenous promoter and as such have physiologically relevant expression levels and spatial patterning of mutant ATXN1 expression. As CAG repeats are unstable, we periodically sequence CAG region to evaluate the number of repeats present in our experimental population. At the time of these experiments, the average number of CAG repeats in our colony was 166. Experimental groups were balanced with an equal number of male and female *Atxn1^154Q/2Q^* and wild-type (WT) mice. In all experiments, investigators were blinded to the genotype.

For fiber photometry experiments, we utilized *Pcp2-Cre;Ai95* mice. *Ai95* line has a floxed-STOP cassette preventing transcription of the *GCaMP6f* fast variant calcium indicator. In *Pcp2-Cre* mice, Cre expression is limited to PCs in the cerebellum, resulting in selective expression of GCaMP6f in PCs in *Pcp2-Cre;Ai95* mice and allowing us to record calcium activity specifically from PCs. *Pcp2-cre;Ai95*mice were crossed with *Atxn1^154Q/2Q^* mice to allow us to record from PCs of SCA1mice.

Animal experimentation was approved by Institutional Animal Care and Use Committee (IACUC) of University of Minnesota and was conducted in accordance with the National Institutes of Health’s (NIH) Principles of Laboratory Animal Care (86–23, revised 1985), and the American Physiological Society’s Guiding Principles in the Use of Animals (National Research Council (US) Committee for the Update of the Guide for the Care and Use of Laboratory Animals, 2011).

### Cerebellar Dissection

Cerebellar anterior and posterior cortex of vermis and hemispheres were dissected as previously described (Hamel and Cvetanovic, 2021). Briefly, brains were placed into a brain matrix with dorsal side up, and one razor blade was placed along the sagittal midline, making sure that the blade pushes all the way to the bottom of the matrix. Six additional razor blades were positioned 1 mm apart from each other (three on each side of the brain). In total there were six sagittal brain sections with the four most lateral sections containing hemispheres where we separated deep cerebellar nuclei from the cortex. Two midline vermal sections were cut just posterior to the primary fissure separating the anterior lobes (I/II-IV/V) from the posterior lobes (VI-X). We used 12 *Atxn1^154Q/2Q^* and wild-type (WT) mice for RNA seq experiments and 8 *Atxn1^154Q/2Q^* and wild-type (WT) mice for RT-qPCR validation. Mice were around 18 weeks of age and balanced for sex.

### RNAsequencing

RNA was extracted from dissected regions using TRIzol Reagent (Life Technologies). RNA was sent to the University of Minnesota Genomics Center for quality control, including fluorometric quantification (RiboGreen assay, Life Technologies) and RNA integrity with capillary electrophoresis (Agilent BioAnalyzer 2100, Agilent Technologies Inc.). All submitted samples with RNA integrity numbers (RINs) 8.0 or greater proceeded to RNA sequencing on an Illumina NextSeq 550 using a 100-nt paired-end read strategy (Friedrich et al., 2018). Data is stored on University of Minnesota Supercomputing Institute Servers. To analyze data raw, paired short reads were analyzed through CHURP (Collection of Hierarchical UMII-RIS Pipelines, PEARC ’19 Proceeding, Article No. 96), which includes data quality control via FastQC, data preprocessing via Trimmomatic (Bolger et al., 2014), mapping via HiSat2 (Kim et al., 2015) against reference mouse genome and expression quantification via Subread (Liao et al., 2014). Resulting count matrix of gene expression was used as input of R (https://www.r-project.org/) package: EdgeR (Chen et al., 2016; Love et al., 2014) (v3.32.1), which tested the differential gene expression and visualized the behavior of expected invariant control genes and published positive control genes. EdgeR was used to reveal sample outliers through Multi-Dimensional Scaling (MDS) and Principle Component Analysis (PCA). Pathway analysis and dot plot creation was preformed using the clusterProfiler package (v3.18.1) using the top 500 DEGs based on absolute value of LogFC (Yu et al., 2012). Heatmaps were created using pheatmap package (https://CRAN.R-project.org/package=pheatmap)(v1.0.12). Volcano plots were created using BiocMananger (https://CRAN.R-project.org/package=BiocManager) (v1.30.16) and EnhancedVolcano (https://github.com/kevinblighe/EnhancedVolcano) (v1.8.0) packages.

### Reverse transcription quantitative polymerase chain reaction (RT-qPCR)

Total RNA was extracted from dissected mouse cerebellar regions using TRIzol (Life Technologies), and RT-qPCR was performed as described previously (Rosa et al., 2022). We used IDT Primetime primers for the *Grid2*, *Gli1,* and *Cck*. GAPDH Universal Probe Library from Roche was used for normalization in the *Cck* experiments. 18S RNA (Forward: AGTCCCTGCCCTTTGTACACA, Reverse: CGATCCGAGGGCCTCACTA) was used for normalization in the *Grid2* and *Gli1* quantification.

### Immunofluorescent (IF) staining

IF was performed on a minimum of four different floating 40-μm-thick brain slices from each mouse (four technical replicates per mouse per region or antibody of interest). Confocal images were acquired using an Olympus FV1000 microscope using an oil 20X objective. Z-stacks consisting of twenty non-overlapping 1-μm-thick slices were taken of each stained brain slice per brain region (i.e., four z-stacks per mouse per region, each taken from a different brain slice). The laser power and detector gain were standardized and fixed between mice within a cohort, and all images for mice within a cohort were acquired in a single imaging session to allow for quantitative comparison. Fluorescent images were acquired using an epifluorescent microscope (Leica DM6000) using a 20X objective. Single 20X images were stitched together to capture one plane of the entire mouse cerebellum. We used 12 mice, *Atxn1^154Q/2Q^* and wild-type (WT), for these experiments, at 18- and 12-weeks of age and balanced for sex.

We used primary antibodies against Purkinje cell marker calbindin (rabbit, Sigma-Aldrich, C9848) as previously described (Asher et al., 2016; Kim et al., 2018). Quantitative analysis was performed using ImageJ (NIH) as described previously. To quantify relative intensity of staining for calbindin we measured the average signal intensity in the region of interest and normalized it to that of the WT littermate control mouse of that cohort. To quantify atrophy of the cerebellar molecular layer we took three measurements per image of the distance from the base of the Purkinje soma to the end of their dendrites, the average being the molecular layer width for that image. We used primary antibodies against vesicular glutamate transporter 2 (VGLUT2) (guinea pig, Millipore, AB2251-I). Extension of climbing fibers was quantified as a ratio of the width of VGLUT2 staining to the width of the molecular layer. We used primary antibodies for astrocytic marker glial fibrillary acidic protein (GFAP) (chicken, Millipore, AB5541). To quantify relative intensity of staining for GFAP we measured the average signal intensity in the region of interest and normalized it to that of the WT littermate mouse. We used primary antibodies for the microglial marker for ionized calcium binding adapting molecule 1 (Iba1) (rabbit, Abcam, AB107159), to quantify the number of microglia in regions of interest.

### Fiber Photometry (Surgery, data acquisition, data analysis)

#### Cannula Implantation

*Atxn1^154Q/2Q^* and wild-type (WT) *Pcp2-Cre;Ai95* mice aged 14 to 15-weeks-old were implanted with a photometry cannula. Briefly mice were induced and anesthetized using isoflurane at 2%. To manage pain, buprenorphine was subcutaneously (SQ) injected 3 hours previous to the surgery, carprofen was delivered SQ right before the start of the surgery, and lidocaine topically applied to the scalp before making the incision. An incision was made to expose the skull, bregma was located, and holes were drilled to implant the fiber photometry cannulas (1.25 mm ferrule, 400um core, 0.37 NA, Neurophotometrics San Diego, CA) into the Lobule 4 (AP: -5.7, ML: -1, and DV: -1.2 from Bregma) and Lobule 6 (AP: -6.6, ML: 1, and DV: -1.4 from Bregma) of the cerebellum. Once the cannula was implanted, it was fixed using dental resin ionomer (Geristore, DenMat). Fiber photometry experiments were not conducted until at least 1 week after the surgery to ensure sufficient time for recovery.

#### Photometry acquisition in Freely Behaving Mice

Fiber photometry recordings took place on a 20-inch by 20-inch arena allowing the mice to move freely for 10 minutes while attached to the patch cord. Using a TDT photometry rig (RZ10X, Tucker-Davis Technologies) the signal from GCamP6f expressed in Purkinje cells was excited and recorded using a 465 nm LED and a 405 nm LED for isosbestic signal sampled at 1 kHz.

#### Fiber photometry visualization and analysis

Raw files from the photometry recording were processed using a custom MATLAB (2022 B) script. This script removes the photobleaching and subtracts the 405 nm signal (isosbestic) from the 465 nm signal (GCaMP6f). Once the file has been processed, OriginPro 2021 (OriginLab) was utilized to normalize the traces to Z-Score. Using the built-in tool Peak Analyzer and the Asymmetric Least Square Smoothing method, the baseline of the trace is determined and artifacts are removed. Calcium events are identified based on the normalized amplitude of the traces with a threshold of 2 standard deviations in 2 second windows. Once the activity peaks were identified for all the traces, they were grouped and frequency as well as amplitude were compared using a one-way ANOVA with Tukey’s post hoc test.

### Statistical analysis

Wherever possible, sample sizes were calculated using power analyses based on the standard deviations from our previous studies, significance level of 5%, and power of 90%. Statistical tests were performed with GraphPad Prism 7.0. Data was analyzed using one-way ANOVA followed by the Tukey’s post-hoc test.

## Data availability

All the data from this study are available from the authors.

## Results

### Increased vulnerability of Purkinje cells in posterior vermis of *Atxn1^154Q/2Q^* mice

To investigate heterogeneity in SCA1 pathology across the cerebellum, we analyzed Purkinje cell (PC) health and glial reactivity across different regions of the cerebellum. This included four lobules of the cerebellar vermis: anterior lobule II, the primary fissure (lobules V/VI), and posterior lobules VII and X (Figure 1A) and four lobules of the hemispheres: Crus I and II, the paramedian, and the copula pyramidis of *Atxn1^154Q/2Q^* mice (Figure 2A). All measurements of cerebellar pathology were performed at 18 weeks of age as *Atxn1^154Q/2Q^* mice at this age have already developed motor deficits but have limited PC loss (Watase et al., 2002).

**Figure 1.**
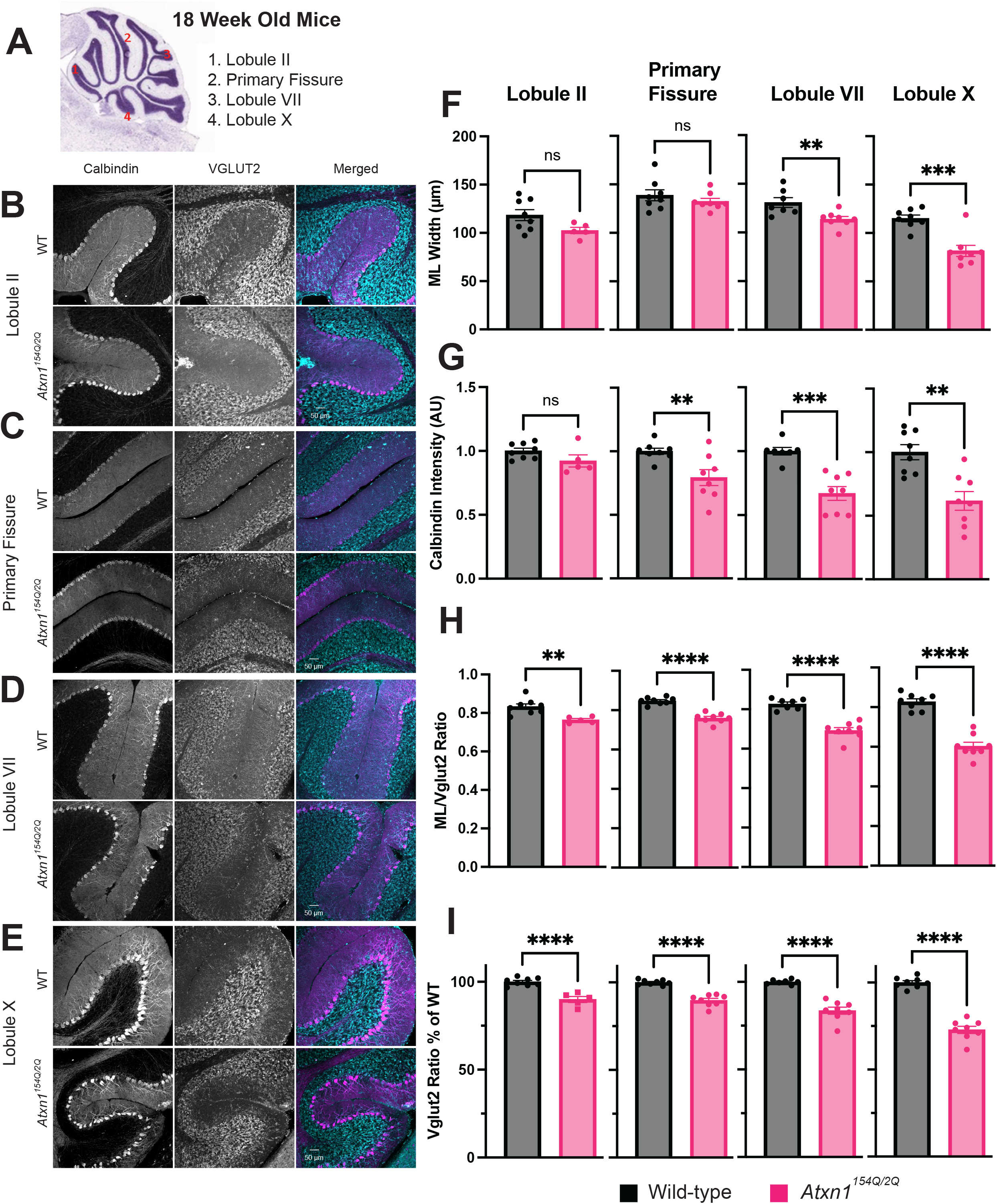
Assessment of dendritic atrophy and synaptic loss in four vermal regions in 18-week-old *Atxn1^154Q/2Q^* mice. **A.** Schematic of sagittal section of mouse cerebellar vermis from Allan Brain Atlas. Lobules II, V/VI, VII, and X were measured and are labeled with red numbers. **B.** Calbindin, vglut2, and merged images of Lobule II for WT (top) and *Atxn1^154Q/2Q^* mice (bottom). **C.** Calbindin, vglut2, and merged images of primary fissure for WT (top) and *Atxn1^154Q/2Q^* mice (bottom). **D.** Calbindin, vglut2, and merged images of Lobule VII for WT (top) and *Atxn1^154Q/2Q^* mice (bottom). **E.** Calbindin, vglut2, and merged images of Lobule X for WT (top) and *Atxn1^154Q/2Q^* mice (bottom). **F.** Quantification of molecular layer thickness of all four vermal lobules. Black bars represent WT, pink bars represent *Atxn1^154Q/2Q^* mice. Student’s t-test, error bars represent SEM, ** = p < 0.01, *** = p < 0.0005. N=8-7 for WT, N=8-5 for *Atxn1^154Q/2Q^* mice. **G.** Quantification of calbindin intensity of all four vermal lobules (arbitrary intensity units). Black bars represent WT, pink bars represent *Atxn1^154Q/2Q^* mice. Students t-test, error bars represent SEM, ** = p < 0.01, *** = p<0.0005. N=8-7 for WT, N=8-5 for *Atxn1^154Q/2Q^* mice. **H.** Quantification of excitatory synapse loss in molecular layer by ratio of VGLUT2 puncta/ molecular layer width measurements in the four vermal lobule regions. Black bars represent WT, pink bars represent *Atxn1^154Q/2Q^* mice. Student’s t-test, error bars represent SEM, ** = p < 0.005, **** = p < 0.0001. N=8-7 for WT, N=8-5 for *Atxn1^154Q/2Q^* mice**. I.** Quantification of percent loss of VGLUT2/ML ratio compared to WT mice. Black bars represent WT, pink bars represent *Atxn1^154Q/2Q^* mice. Student’s t-test, error bars represent SEM, **** = p < 0.0001. N=8-7 for WT, N=8-5 for *Atxn1^154Q/2Q^* mice.

**Figure 2.**
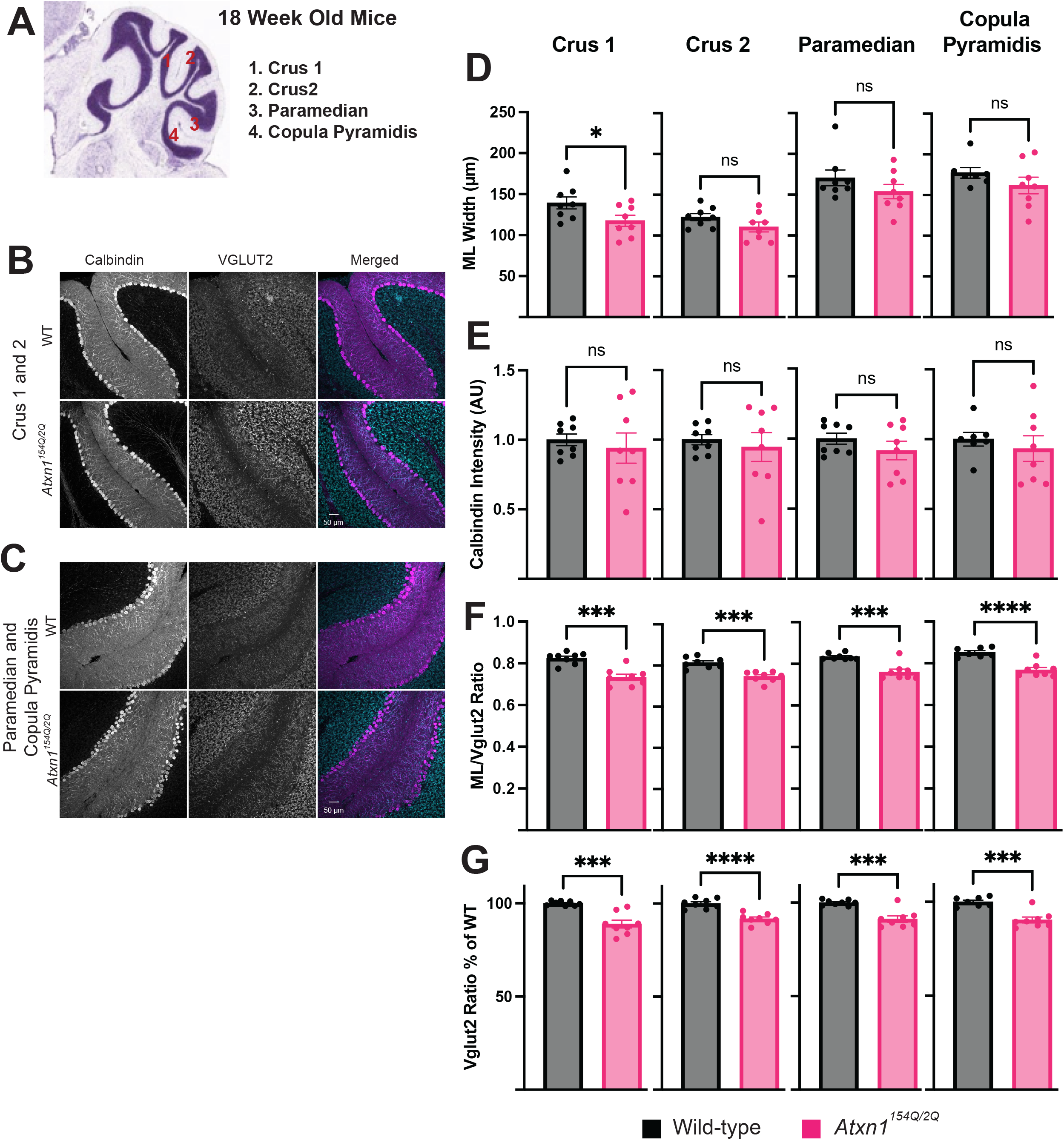
Assessment of dendritic atrophy and synaptic loss in four hemisphere regions in 18-week-old *Atxn1^154Q/2Q^* mice. **A.** Schematic of sagittal section of mouse cerebellar hemispheres from Allan Brain Atlas. Crus 1, crus 2, paramedian, and copula pyramidis were measured and are labeled with red numbers. **B.** Calbindin, vglut2, and merged images of Crus 1 and 2 for WT (top) and *Atxn1^154Q/2Q^* mice (bottom). **C.** Calbindin, vglut2, and merged images of paramedian and copula pryamidis WT (top) and *Atxn1^154Q/2Q^* mice (bottom). **D.** Quantification of molecular layer thickness of all four hemisphere lobules. Black bars represent WT, pink bars represent *Atxn1^154Q/2Q^* mice. Student’s t-test, error bars represent SEM, * = p < 0.05. N=8-6 for WT, N=8-7 for *Atxn1^154Q/2Q^* mice. **E.** Quantification of calbindin intensity of all four hemisphere lobules (arbitrary intensity units). Black bars represent WT, pink bars represent *Atxn1^154Q/2Q^* mice. Students t-test, error bars represent SEM. N=8-7 for WT, N=8-7 for *Atxn1^154Q/2Q^* mice. **F.** Quantification of excitatory synapse loss in molecular layer by ratio of VGLUT2 puncta/ molecular layer width measurements in the four hemisphere lobule regions. Black bars represent WT, pink bars represent *Atxn1^154Q/2Q^* mice. Student’s t-test, error bars represent SEM, *** = p < 0.0005, **** = p < 0.0001. N=8-7 for WT, N=8-7 for *Atxn1^154Q/2Q^* mice**. G.** Quantification of percent loss of VGLUT2/ML ratio compared to WT mice. Black bars represent WT, pink bars represent *Atxn1^154Q/2Q^* mice. Student’s t-test, error bars represent SEM, *** = p < 0.0005, **** = p < 0.0001. N=8-7 for WT, N=8-7 for *Atxn1^154Q/2Q^* mice.

PCs pathology in this and other SCA1 mouse models is quantified by the atrophy of PC dendrites, reduced calbindin (a calcium binding protein that is highly expressed in PCs) expression, and loss of vesicular glutamate transporter 2 positive (VGLUT2+) climbing fiber synapses on PC dendrites (Ebner et al., 2013; Kim et al., 2018). To compare these changes across the cerebellum, we stained sagittal cerebellar sections with the PC marker calbindin and VGLUT2 (Figure 1B-E and 2B-C). We only found significant dendritic atrophy of PCs, measured as reduced width of the molecular layer (ML) in posterior lobules VII and X of the vermis (Figure 1F) and in Crus 1 of the hemispheres (Figure 2D). Calbindin intensity was significantly decreased only in the posterior vermis-specifically lobules VI, VII and X (Figure 1G and Figure 2E).

While PC dendritic atrophy and reduced calbindin expression were largely limited to posterior cerebellar vermis, a significant loss of climbing fiber VGLUT2 synapses on PC dendrites was detected in all lobules examined (Figure 1H and Figure 2F). Notably, this loss was graded along the anterior to posterior axis with lobule II, primary fissure and those in the hemispheres having a 10% decrease, lobule VII having a 30% decrease, and most posterior lobule X having the largest (a 40% reduction) synaptic loss (Figure 1I and Figure 2G).

We also measured the ML thickness of each lobule in the cerebellum, normalized to the wild-type lobules, and combined the measurements based on region (Supplementary Figure 1A). We found a significant reduction in the ML width in the posterior lobules of the vermis (Lobules VI-X) and in the hemispheres (Supplementary Figure 1B) of *Atxn1^154Q/2Q^* mice but not in the anterior vermal lobules (Lobules I-V).

We investigated regional pathology at 12 weeks of age, around the onset of motor deficits, to determine whether these observed regional differences are present at an earlier disease stage. Notably, we found significant reduction in calbindin intensity only in the posterior lobules VII and X (Supplementary Figure 2D). When assessing the loss of climbing fiber synapses, we found a statistically significant reduction in the lobules V/VI, VII, and X (Supplementary Figure 2C and E). Moreover, anterior lobule II showed no significant changes in any of our PC pathology measures at the 12-week time point.

These results show that PC pathology presents earliest and most severely in the posterior vermis. Moreover, there is relative sparing of PC degeneration in the anterior vermis. This is consistent with the increased vulnerability of posterior cerebellar PCs observed in human SCA1 patients.

### Glial pathology is markedly severe in posterior SCA1 vermis

To examine intracerebellar differences in SCA1 glial pathology we evaluated expression of glial fibrillary acidic protein (GFAP) and the density of Iba1 labeled microglia as a simple measure of reactive astrogliosis and microgliosis. Bergmann glia (BG) reactivity can be quantified as an increase in the expression of GFAP in the molecular and Purkinje cell layers, and was previously described in the cerebellum of SCA1 mouse models (Kim et al., 2018; Rosa et al., 2022). While there are many different types of glia in the cerebellum, we chose to focus on BGs because of their close association with PCs in the molecular layer. BG processes closely interact with PC dendrites and are critical to ongoing PC health and function (Bellamy, 2006; Farmer et al., 2016; Luttik et al., 2022). We measured GFAP immunofluorescence using confocal microscopy (Figure 3A and B) and found a significant increase in BG GFAP intensity only in Lobule X of *Atxn1^154Q/2Q^* mice, indicating selective astrocytic reactivity in lobule X in SCA1. Interestingly, the BG of wild-type mice also exhibit higher GFAP expression in lobule X compared to other vermal lobules (Figure 3C). This suggests that there is an intrinsic regional heterogeneity of BG in the cerebellum. Finally, as with BGs, microglial density was increased in vermal lobule X (Figure 3D) and Crus 1 of the hemispheres (Figure 3E) in SCA1 mice compared to controls.

**Figure 3.**
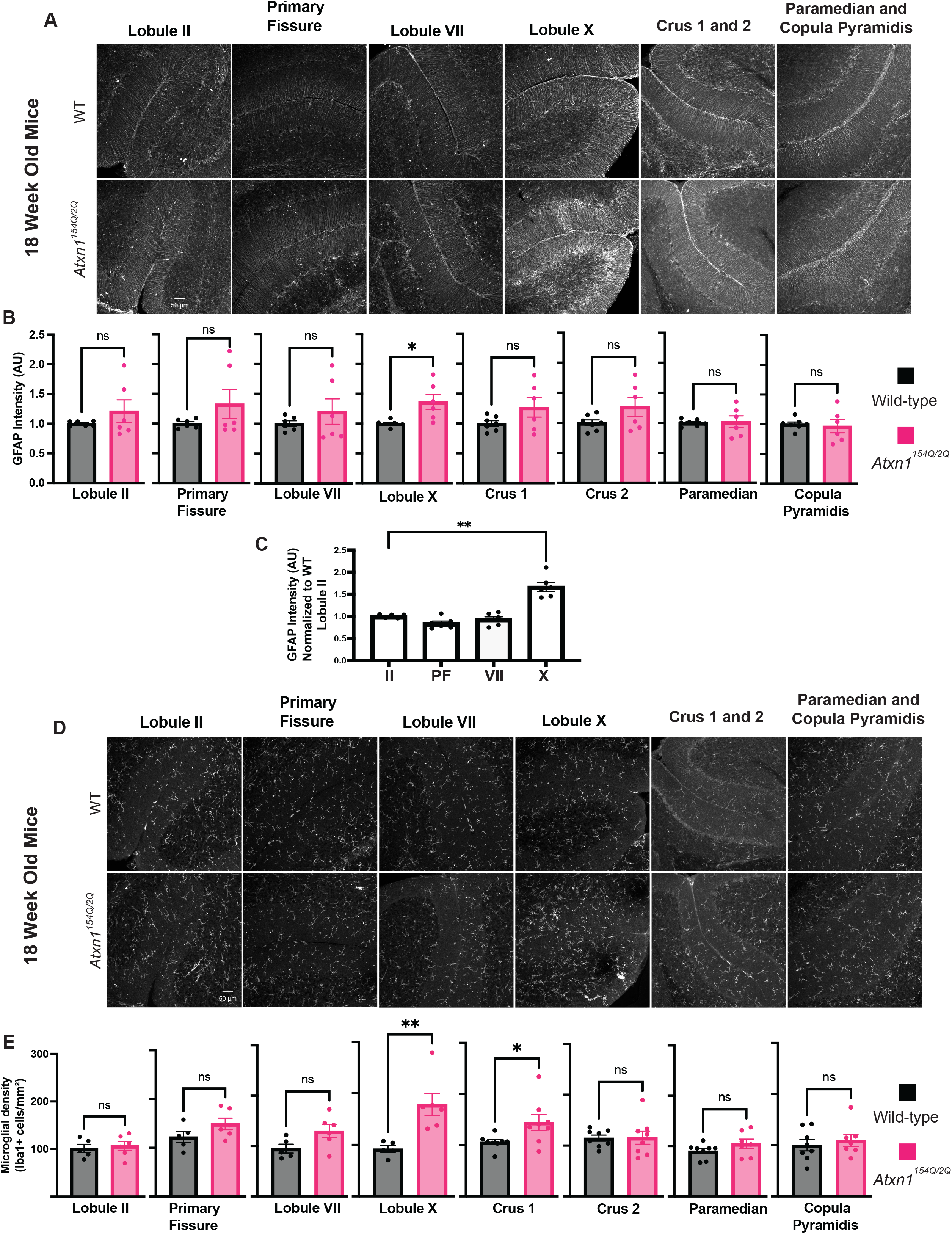
Assessment of gliosis across four regions of the cerebellar vermis and four regions of cerebellar hemispheres 18-week-old *Atxn1^154Q/2Q^* mice. **A.** Images of GFAP staining in the cerebellar vermis lobules II, primary fissure, VII, and X, and hemisphere lobules crus1, crus1, paramedian, and copula pyramidis. WT on top, *Atxn1^154Q/2Q^* mice (bottom). **B.** Quantification of intensity of GFAP signal for all eight regions of the cerebellum, measured in arbitrary intensity units (AUI). Black bars represent WT, pink bars represent *Atxn1^154Q/2Q^* mice. Student’s t-test, error bars represent SEM, * = p < 0.01. N=8-6 for WT, N=8-5 for *Atxn1^154Q/2Q^* mice. **C.** Quantification of intensity differences across the four cerebellar vermis regions in wild-type mice. Intensity measurements normalized to Lobule II. One-way ANOVA with Bonferroni post-hoc test, error bars represent SEM, ** = p < 0.005, N=6. **D**. Images of Iba1 staining in the in the cerebellar vermis lobules II, primary fissure, VII, and X, and hemisphere lobules crus1, crus1, paramedian, and copula pyramidis. WT on top, *Atxn1^154Q/2Q^* mice (bottom). **E.** Quantification of density of Iba1 positive cells (microglia) in the molecular layer for all eight regions of the cerebellum, measured in arbitrary intensity units (AUI). Black bars represent WT, pink bars represent *Atxn1^154Q/2Q^* mice. Student’s t-test, error bars represent SEM, * = p < 0.05, ** = p < 0.01. N=8-6 for WT, N=8-5 for *Atxn1^154Q/2Q^* mice.

Assessment of gliosis at the “early disease” 12 week time point demonstrated increased GFAP staining intensity only in lobule X (Supplementary Figure 2F and G) and a trending increase in microglial density in lobule X of the 12-week *Atxn1^154Q/2Q^* mice (Supplementary Figure 2H and I).

Taken together, we show that in early stages of disease, the first signs of pathology arise in the posterior vermis, including both evidence of PC degeneration and activation of BG. During moderate stages of pathology, both PCs and glia exhibit pathology that is markedly severe in the posterior vermis.

### Transcriptional heterogeneity across cerebellar regions

We reasoned that a transcriptomic study of cerebellar regions could provide crucial insight into cerebellar heterogeneity and how it contributes to and is impacted by cerebellar disease.. We employed a quick and reproducible dissection method that we developed and validated (Hamel and Cvetanovic, 2021) to isolate distinct cortical regions of the cerebellum-cerebellar cortex of the anterior vermis (CCAV), cerebellar cortex of the posterior vermis (CCPV), and cerebellar cortex of the hemispheres (CCH)-from *Atxn1^154Q/2Q^* mice and wild-type littermate controls (Supplementary Figure 3A).

Three different analytical paradigms were used to determine the regional heterogeneity of the cerebellum in different contexts. First, we performed a comparative analysis across regions in wild-type animals to identify the unique genes and pathways enriched in each cerebellar region (Supplementary Figure 3B, yellow arrows). Then, to provide insight into molecular mechanisms underlying selective vulnerability in SCA1, we compared transcriptomes of anterior vermis, posterior vermis and hemispheres in *Atxn1^154Q/2Q^* mice to wild-type controls (Supplementary Figure 3B, red arrows). Finally, we directly compared between our isolated cerebellar regions in SCA1 mice to investigate how cerebellar heterogeneity is impacted in disease.

### Compelling cerebellar heterogeneity of gene expression in wild-type mice

We compared gene expression across CCAV, CCPV, CCH) tin wild-type mice (Supplementary Figure 3B, yellow arrows) and ranked the top 15 differentially expressed genes (up and down regulated) from each comparison (Figure 4A-C). While this shows that each region has unique molecular characteristics, it does not capture the magnitude of these differences across regions. Intriguingly, we found that the anterior vermis of the cerebellum is notably distinct from both the posterior vermis and the lateral hemispheres. A total of 1229 differentially expressed genes (DEGs) were changed across the anterior to posterior axis of the vermis, with 884 genes (71.9%) increased in expression in the anterior vermis (Figure 4D). Comparison of the anterior vermis to the lateral hemispheres demonstrated a similar magnitude of transcriptomic uniqueness with 77.9% of the 774 identified DEGs upregulated in the anterior vermis (Figure 4E). This could, in part, be due to a degree of molecular similarity between the posterior vermis and lateral hemispheres, as there were only 365 DEGs identified between these two regions (Figure 4F).

**Figure 4.**
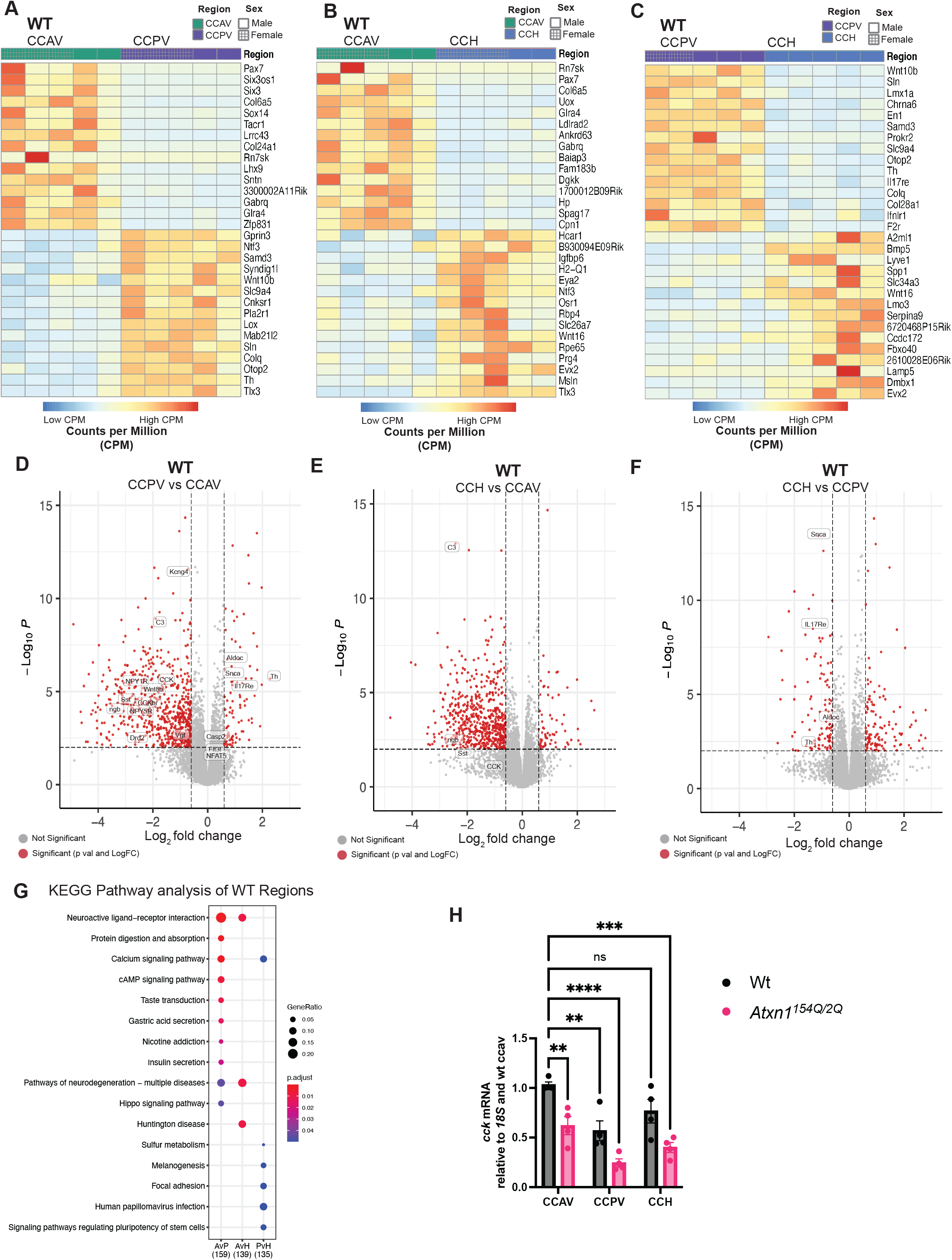
Transcriptomic analysis in wild-type mice identifies differently regulated genes across the cerebellar cortex. **A-C.** Heatmaps of top up and down regulated DEGs in between the anterior and posterior vermis (A), the anterior vermis and the cerebellar hemispheres (B), and the posterior vermis and cerebellar hemispheres (C). Heatmaps were generated by ranking the top 15 up and down regulated DEGs based on logFC value and plotting the counts per million (CPM) of each gene in individual samples. Expression is normalized per gene in the heatmap representation. **D-F.** Volcano plots of all DEGs between the anterior and posterior vermis (D), the anterior vermis and the cerebellar hemispheres (E), and the posterior vermis and cerebellar hemispheres (F). Red dots represent significant DEGs based on both p-value and logFC, gray dots represent DEGs significant by one metric or that did not reach significance. **G.** KEGG pathway analysis of enriched pathways of top 500 DEGs between the anterior and posterior vermis, the anterior vermis and the cerebellar hemispheres, and the posterior vermis and cerebellar hemispheres. Dot size represents gene ratio (number of genes in the enriched pathway divided by the total number of DEGs), and color of the dot represents p-value. **H.** *Cck* mRNA expression across cerebellar cortical regions in WT and *Atxn1^154Q/2Q^* mice by RTqPCR. WT in black bars, *Atxn1^154Q/2Q^* in pink bars. n = 4 per genotype. Error bars = SEM, Two-way ANOVA with Bonferroni’s multiple correction. ** p < 0.005, *** p <0.001, **** p <0.0001.

To understand how these differentially expressed genes could be functionally contributing to regional vulnerability, we assessed pathways altered using KEGG pathway analysis. We identified the top ten pathways that are enriched in the comparison between anterior and posterior vermis, including neuroactive ligand, protein digestion, calcium, cAMP, and neurodegeneration (Figure 4G). Interestingly, several of these pathways have been implicated in SCA1. For instance, *Cck* and *Cckbr*, which encode for neuroactive ligand Cholesytokinin (CCK) and its receptor, were recently found to be neuroprotective in SCA1 and SCA2 (Ingram et al., 2016; Wozniak et al., 2021). We confirmed increased *Cck* expression in the anterior vermis by RT-qPCR (Figure 4H). Additionally, the anterior vermis is enriched in genes negatively regulating neuroinflammation including anti-inflammatory neuropeptide Y (NPY),its receptors 1 and 5 (NPY1R and NPY5R) (Li et al., 2019), and dopamine 2 receptor (*DRD2*) (Du et al., 2018; Zhu et al., 2020) (Figure 4D). Increased expression of anti-inflammatory genes could contribute to reduced pathology in the anterior vermis of SCA1 mice.

Conversely, increased expression of inflammatory regulating inflammation (*IL17Re* (*Interleukin 17 Receptor*), *NFAT* 5 *(nuclear factor of activated T cells 5*) and *Sp1)*, cell death (*Cas2* (c*aspase-2*)), and dysregulated synapse (*Snca* (α*Synuclein*)) genes in the healthy posterior vermis could predispose/contribute to the selective vulnerability observed in SCA1 (Figure 4E and F).

### Molecular differences in SCA1 pathogenesis across cerebellar cortex

Our next goal was to provide insight into molecular mechanisms of disease pathogenesis in SCA1. We achieved this by investigating DEGs in anterior vermis, posterior vermis, and hemispheres of *Atxn1^154Q/2Q^* mice compared to their wild-type littermate controls. A similar number of DEGs were identified in each of these regions (CCAV: 1655, CCPV: 1715 and CCH: 1445) (Figure 5A). Consistent with previous findings from whole cerebellar RNA sequencing (Driessen et al., 2018; Ingram et al., 2016), the majority of the DEGs identified in each region were downregulated in the *Atxn1^154Q/2Q^* mice (Figure 5B).

**Figure 5.**
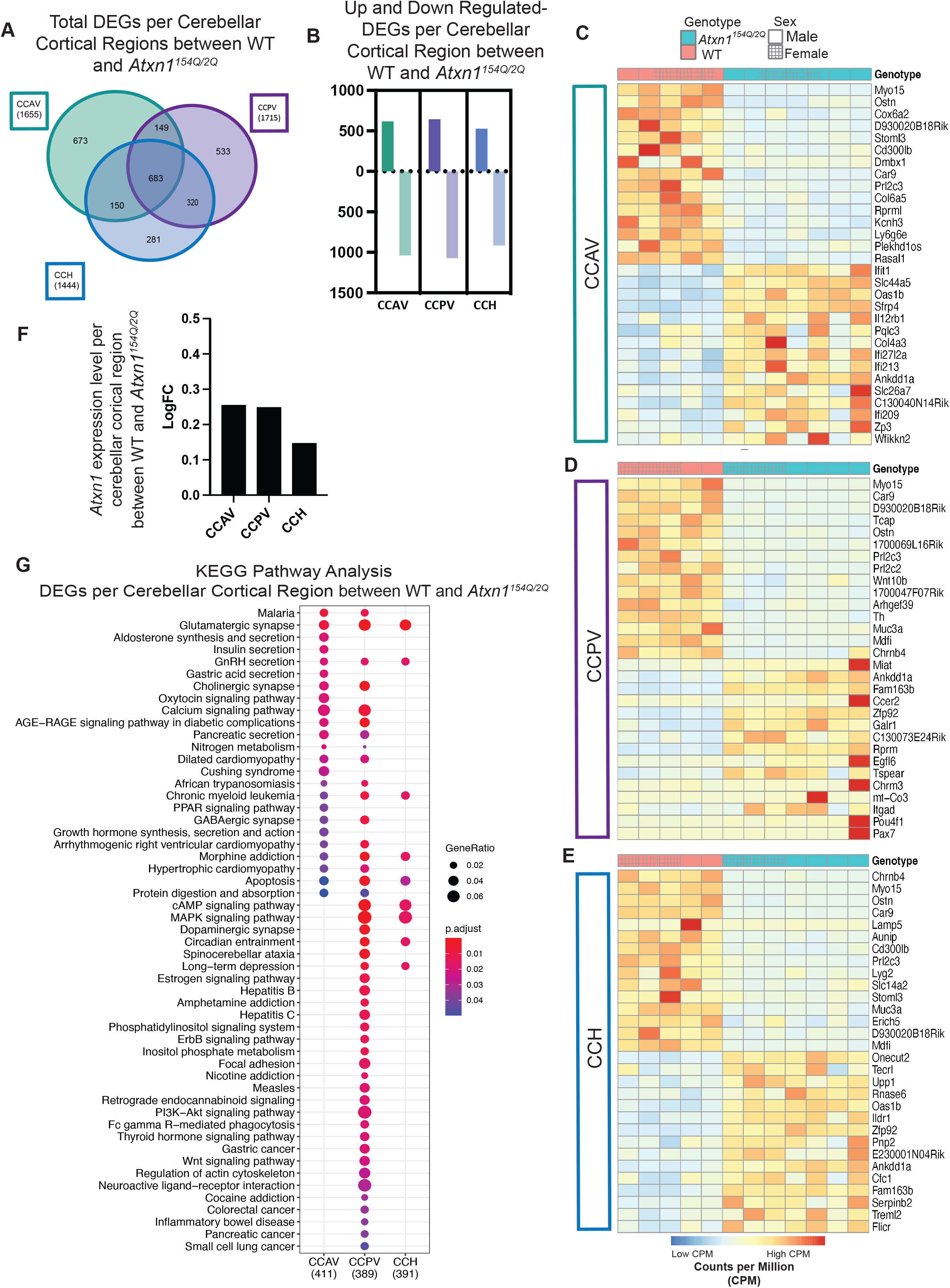
SCA1 intracerebellar transcriptomics identifies shared and unique disease pathways in *Atxn1^154Q/2Q^* mice. **A.** Venn diagram of DEGs in each cerebellar cortical region. 1655 DEGs in CCAV (green), 1755 DEGs in CCPV (purple), and 1444 DEGs in CCH (blue). **B.** Number of up and down regulated DEGs in CCAV, CCPV, and CCH. **C-E.** Heatmaps of top up and down regulated DEGs in between WT and *Atxn1^154Q/2Q^* mice in each cerebellar cortical region: CCAV (C), CCPV (D), and CCH (E). Each column represents an individual sample, WT in pink, *Atxn1^154Q/2Q^* in teal, females with hashed boxes, males with solid boxes. Heatmaps were generated by ranking the top 15 up and down regulated DEGs based on logFC value and plotting the counts per million (cpm) of each gene in individual samples. Expression is normalized per gene in the heat map representation. **F.** KEGG pathway analysis of enriched pathways of top 500 DEGs between WT and *Atxn1^154Q/2Q^* mice in each cerebellar cortical region. Dot size represents number of genes in the enriched pathway, and color of the dot represents p-value.

Approximately 42% (683) DEGs were altered in all three regions of interest, constituting a shared effect of SCA1 pathogenesis. Although region selective pathology is apparent in *Atxn1^154Q/2Q^* mice at this time point, we did not find that the expression of this shared gene set was any more pronounced in the posterior vermis. In keeping with this high proportion of shared gene expression changes across regions, Gene Ontology (GO) pathway analysis found that pathways regulating of ion transmembrane transport, voltage gated channel activity, and potassium ion transmembrane transport were consistently changes across regions. Notably, many of these pathways have been identified in bulk cerebellar transcriptomic studies and have been functionally validated in physiological analysis of SCA1 cerebella (citations for both). (Supplementary Figure 4A and B).

While a large proportion of our transcriptomic data is consistent with and supported by previous bulk analysis of the SCA1 cerebellum, we also found a surprisingly large number of unique differential gene expression changes in each cerebellar region of SCA1 mice. In fact, most of the DEGs with the greatest log fold change (LogFC) were unique to each region, suggesting distinct differences in disease pathogenesis across cerebellar regions. Importantly, *Atxn1* expression did not differ across regions, indicating that regional differences in SCA1 pathogenesis across cerebellum are not a consequence of differences in *Atxn1* expression (Figure 5F). Differentially expressed genes that are unique to each of the cerebellar cortical regions could reflect and contribute to region specific pathology (Figure 5C-E). For instance, *Grid2* is significantly decreased in the posterior vermis and hemispheres but not in the anterior cerebellum (Figure 6C). *Grid2* (Glutamate Ionotropic Receptor Delta Type Subunit) encodes a glutamate ionotropic receptor GRID2 which is highly expressed in PCs. Loss of GRID2 expression through homozygous deletion causes spinocerebellar ataxia type 18 (Sarna and Hawkes, 2003; Selimi et al., 2003)(Hills et al., 2013; Utine et al., 2013). We confirmed the region-selective decrease of *Grid2* expression in the posterior vermis using an independent set of RNA isolated from these three regions and RTqPCR (Figure 6D). Following our observation that glial reactivity was enhanced in the posterior vermis, we also found reduced expression of genes important for BG function in the posterior vermis and hemispheres. This includes a notable downregulation of *Kcnj10* and *Slc1a3* (Figure 6B); these genes are responsible for regulating homeostasis of potassium and glutamate (Djukic et al., 2007; Rothstein et al., 1996), respectively. Both of these genes have been previously shown to be downstream of Sonic hedgehog (Shh) signaling in the cerebellum (Farmer et al., 2016). Examination of Shh receptor (*Ptch1 and Ptch2)* and downstream transcription factor (*Gli1*) expression in this system showed that the posterior cerebellum had the most severe decrease in the expression (Figure 6E). The enhanced decrease of *Gli1* was confirmed with RT-qPCR (Figure 6F). Together, these results indicate that pronounced impairment of Shh signaling in the posterior cerebellum may be contributing to the regionally worsened BG dysfunction in SCA1.

**Figure 6.**
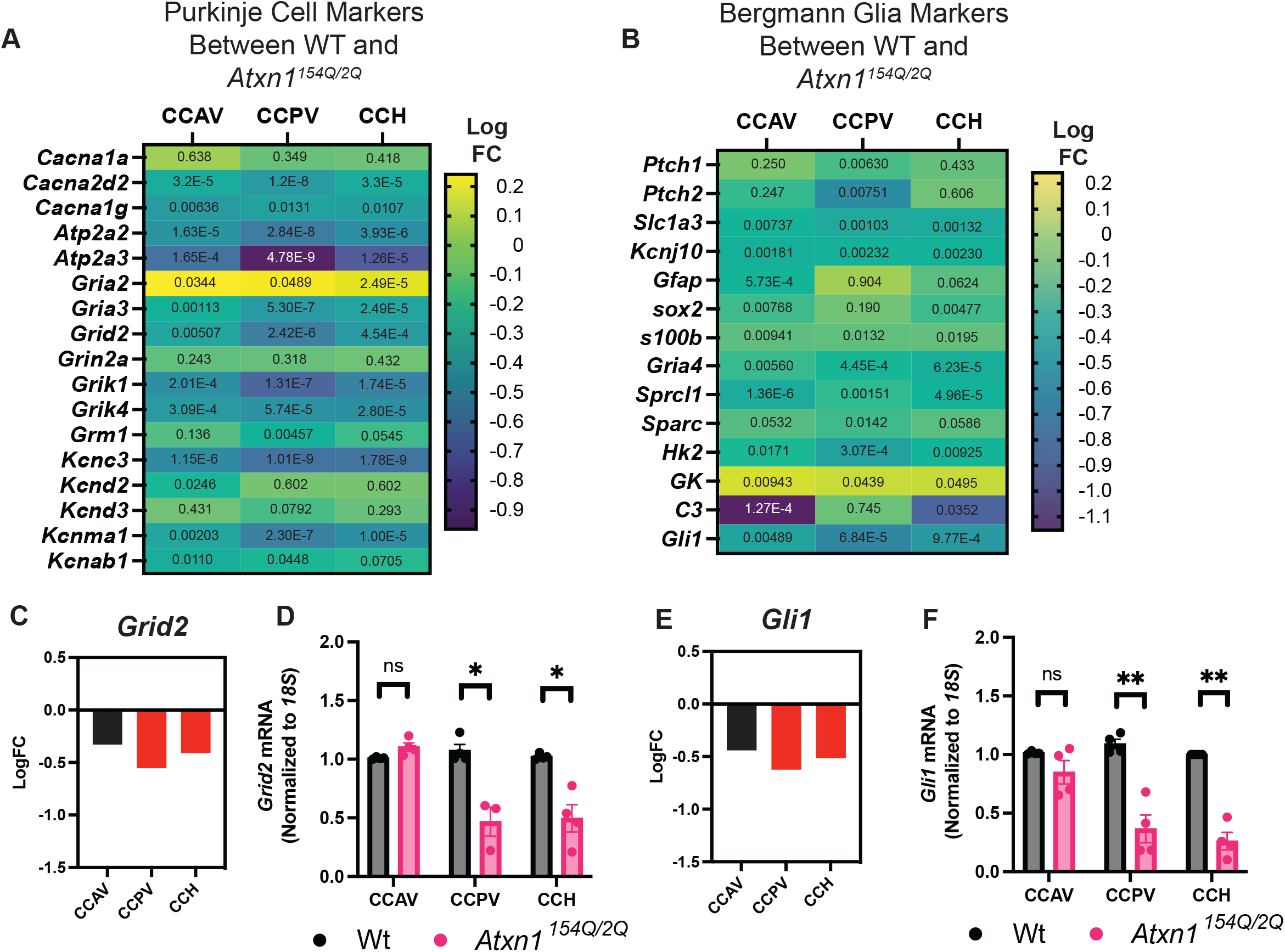
Cell-specific gene expression across the cerebellar cortical regions. **A.** Heatmap of Purkinje cell specific signaling gene expression differences in *Atxn1^154Q/2Q^* mice based on logFC across each cerebellar cortical region; anterior vermis (left), posterior vermis (middle), cerebellar hemispheres (right). **B.** Heatmap of Bergmann glia specific gene expression differences in *Atxn1^154Q/2Q^* mice based on logFC across each cerebellar cortical region; anterior vermis (left), posterior vermis (middle), cerebellar hemispheres (right). P values are listed in for each gene in each region. **C.** *Grid2* expression from RNAsequencing results across cerebellar cortical regions; red bars mark significance based on FDR < 0.05, black bars mark lack of significance based on FDR > 0.05. **D.** *Grid2* mRNA expression across cerebellar cortical regions in WT and *Atxn1^154Q/2Q^* mice by RTqPCR. WT in black bars, *Atxn1^154Q/2Q^* in pink bars. n = 4 per genotype. Error bars = SEM, Welch’s t-test, * p < 0.05. **E.** *Gli1* expression from RNAsequencing results across cerebellar cortical regions; red bars mark significance based on FDR < 0.05, black bars mark lack of significance based on FDR > 0.05. **F.** *Gli1* mRNA expression across cerebellar cortical regions in WT and *Atxn1^154Q/2Q^* mice by RTqPCR. WT in black bars, *Atxn1^154Q/2Q^* in pink bars. n = 4 per genotype. Error bars = SEM, Welch’s t-test, * p < 0.05, ** p<0.005.

To KEGG pathway analysis identified difference is the pathways disrupted across the regions. Notably, the largest number of disrupted pathways was in the posterior cerebellum (Figure 5G). This result may indicate that number of altered pathways may be a significant contributor to selective vulnerability.

### SCA1 reduces intrinsic intracerebellar heterogeneity

Finally, we asked how SCA1 impacts intrinsic cerebellar regional heterogeneity by comparing gene expression across cerebellar regions in *Atxn1^154Q/2Q^* mice (Supplementary Figure 3B, pink arrows). Based on the number of differentially expressed genes found per comparison, the regional distinctions between the anterior and posterior vermis shrank while the hemispheres remained transcriptionally unique. In contrast to the 1229 genes that were differentially expressed between the anterior and posterior vermis in the wild-type mice, this comparison yielded only 507 DEGs in *Atxn1^154Q/2Q^* mice (Figure 7A). Moreover, the number of region-unique DEGs dropped from 671 genes to 186 in *Atxn1^154Q/2Q^* mice in the comparison of the anterior and posterior vermis (Figure 7B). Similarly, the number of DEGs between posterior vermis and hemispheres was slightly decreased (from 365 DEGs in wild-type mice to 213 in *Atxn1^154Q/2Q^* mice (Figure 7B). Intriguingly, this trend did not extend to the hemispheres. The number of DEGs identified between the SCA1 anterior vermis and hemispheres was 705, similar to 744 DEGs found in wild-type mice. This large reduction of number of DEGs between posterior and anterior vermis or hemispheres suggests significant loss of heterogeneity in the *Atxn1^154Q/2Q^* mice.

**Figure 7.**
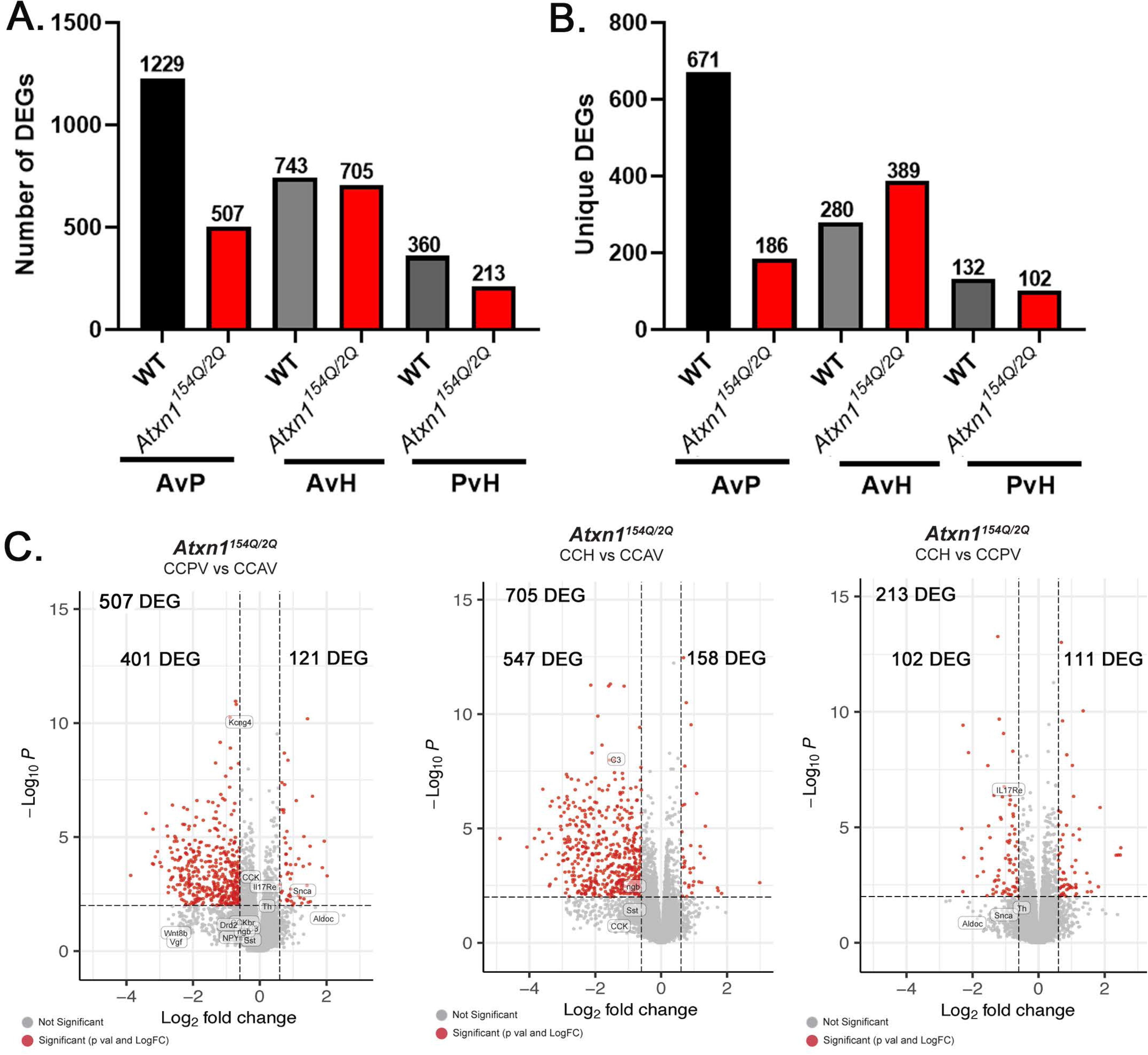
Reduced heterogeneity in SCA1 cerebellar regions. **A.-B.** Transcriptomic analysis in wild-type and in SCA1 mice identifies differently regulated genes across the cerebellar cortex and genes uniquely expressed in either of the regions in pairwise comparison. **C.** Volcano plots of all DEGs between the anterior and posterior vermis (left), the anterior vermis and the cerebellar hemispheres (middle), and the posterior vermis and cerebellar hemispheres (right). Red dots represent significant DEGs based on both p-value and logFC, gray dots represent DEGs signific5ant by one metric or that did not reach significance.

Comparing the top 15 differentially expressed genes (up and down regulated) from each region further reveals impact of SCA1 on intrinsic heterogeneity (Supplementary Figure 5A and Figure 4). Furthermore, out of the top ten pathways identified by KEGG pathway analysis differing anterior and posterior vermis in WT mice (Figure 4G), only one (Hippo signaling) remained different between anterior and posterior vermis in SCA1 mice (Supplementary Figure 5D, Figure 7D). These results indicate that SCA1 profoundly alters intrinsic heterogeneity of gene expression between anterior and posterior vermis.

### Regional heterogeneity of PC activity during free unrestricted walking and non-walking states in wild-type and SCA1 mice

Previous studies demonstrated intrinsic differences in PC activity across the cerebellum. In particular most of PC in the posterior cerebellum were marked by zebrin expression and exhibited a lower firing rate compared to zebrin negative PCs in the anterior cerebellum (Zhou et al., 2014). In *Atxn1^154Q/2Q^* mice, cerebellar slice electrophysiology revealed reduced regularity of PC firing rate, measured as increased coefficient of variation of the inter-spike interval (Bushart et al., 2021).Seeking to build off of this previous work, we sought to investigate regional PC activity *in vivo* during cerebellum regulated behavior such as self-motivated and unrestrained walking. We then investigated whether the regional pattern of PC activity is altered in disease and whether activity of posterior PC is more impacted in SCA1 mice.

We quantified PC activity across the anterior to posterior axis of the cerebellar vermis in freely moving mice by employing fiber photometry and PC-specific expression of the genetically encoded calcium indicator (GCAMP6f). Fiber photometry is a relatively less invasive surgery than miniscopes, uses lightweight instrumentation that minimizes effects on balance, and it allows for long-term monitoring of PC activity in freely moving mice. Specific expression of GCaMP6f in PCs was achieved by crossing the *Ai95* line, which has a floxed-STOP cassette preventing transcription of the GCaMP6 fast variant calcium indicator, with *Pcp2-Cre* which expresses Cre recombinase only in Purkinje cells (Madisen et al., 2015; Sługocka et al., 2017)(Chen et al., 2013). Fiber photometry cannulas were implanted into anterior lobule IV and the contralateral posterior lobule VI (Figure 8A) which are as far anterior and posterior we could access while minimizing the damage to the cerebellum and surrounding tissue. Post-surgery, mice were allowed to recover for one week after which we recorded mouse movement along with the calcium activity during a ten minute period of unrestricted, self-motivated walking and not walking states in the open field (Figure 8).

**Figure 8.**
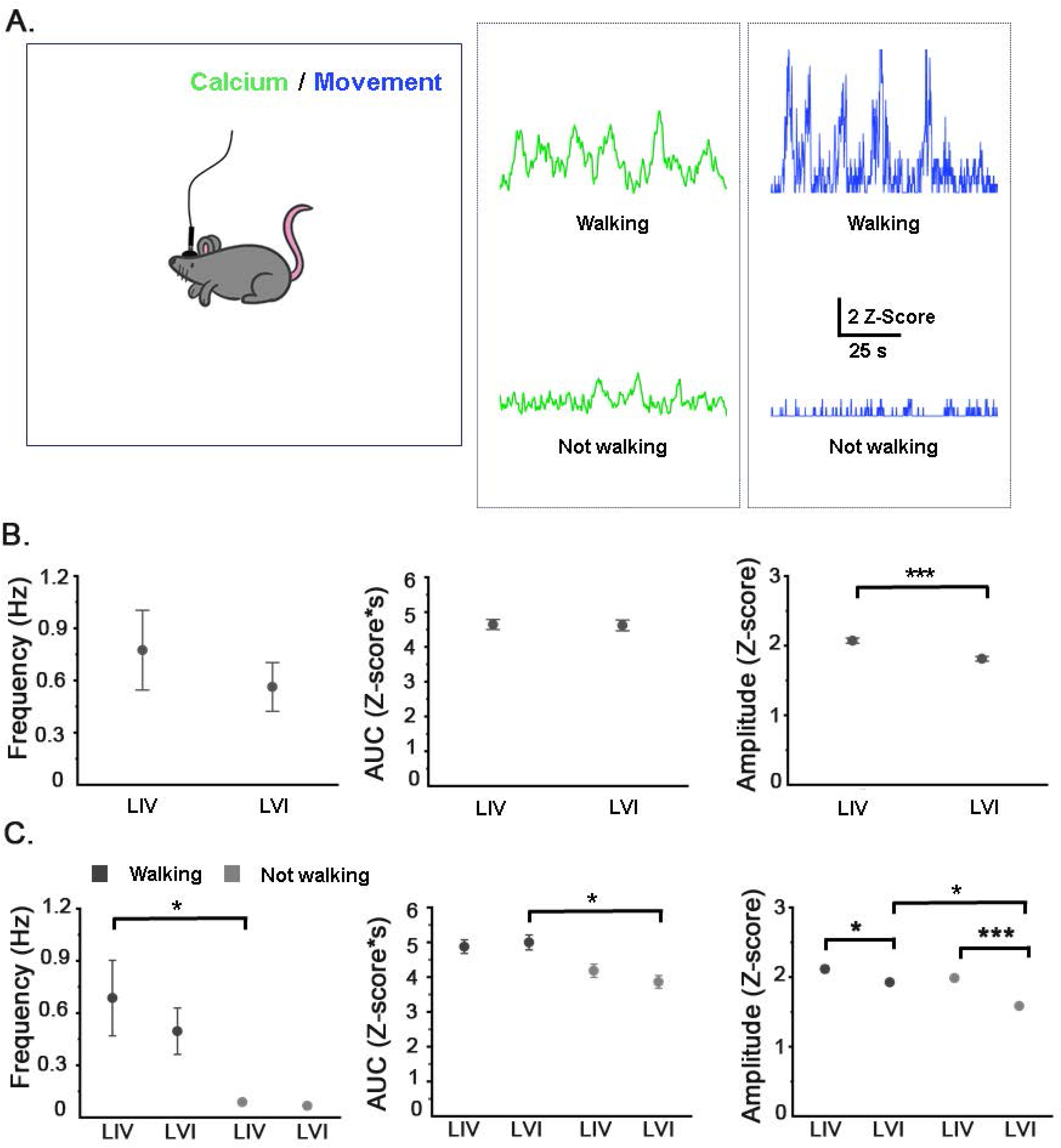
PCs activity in the anterior and posterior vermis. **A.** Schematics of experiment and example traces of calcium activity (green) and movement (blue) **B.** Frequency, amplitude and area under the curve (AUC) of PCs calcium events over ten minute exploration in the open field in lobules IV (LIV) and VI (LVI) of wild-type mice. **C.** Frequency, amplitude and area under the curve (AUC) of PCs calcium events during walking (black circles) and non-walking states (grey circles) in lobules IV (LIV) and VI (LVI) of wild-type mice. N = 5 wild-type mice 16-18 weeks of age. One-way ANOVA with Tukey’s post hoc * p< 0.05, *** p < 0.001.

Overall, the amplitude of PC calcium events was significantly larger in the anterior IV compared to posterior lobule VI of wild-type mice (Figure 8). The frequency of calcium events was also slightly larger in the lobule IV, but it did not reach statistical significance (Figure 8). Together, these results support previous work suggesting that areas rich in zebrin+ PCs should have reduced firing strength. Further, it suggests that the cerebellar regional heterogeneity occurs both at a transcriptional and a functional level.

The cerebellum is particularly important for the execution of coordinated movements and maintaining balance while walking. Notably, SCA1 patients exhibit difficulties in balance and coordinated movement while walking. Therefore, we next analyzed the data considering the state of movement to determine PC activity in lobules IV and VI during walking and non-walking states. We found that PC activity is increased during walking in both lobules, though the manner to which this changes during locomotion is region-specific. Walking drives an increase in frequency in lobule IV while in lobule VI there is an increase in the amplitude and area under the curve (AUC) when compared to rest (Figure 8C). This result indicates that walking correlates with increased activity of PCs across the cerebellum, but using a different mechanisms.

We next examined PC activity in lobules IV and VI in *Atxn1^154Q/2Q^* mice. We found decreased peak amplitude in both of lobules of *Atxn1^154Q/2Q^* mice compared to WT littermate controls (Figure 9A-B top row). In lobule IV we also found a trending decrease in the AUC (P = 0.053), while in lobule VI there was a trending increase both in frequency and AUC (p= 0.054 and 0.08 respectively). This suggests differences in the manner through which PC activity is altered in these lobules in SCA1.

**Figure 9.**
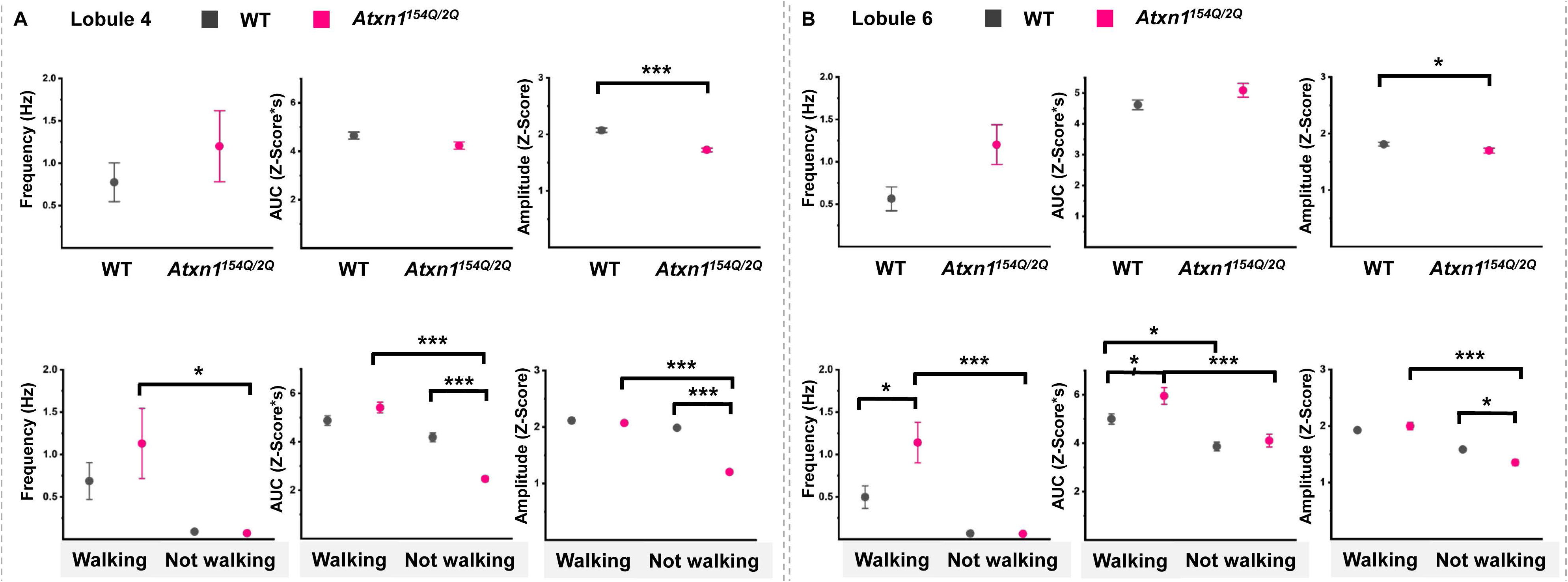
PCs activity in *Atxn1^154Q/2Q^* mice. **A.** Frequency, amplitude and area under the curve (AUC) of PCs calcium events in lobule IV (LIV) of wild-type and *Atxn1^154Q/2Q^* mice over ten minute exploration in the open field (top) or during walking (black circles) and non-walking states (grey circles) (bottom). **B.** Frequency, amplitude and area under the curve (AUC) of PCs calcium events in lobule VI (LVI) of wild-type and *Atxn1^154Q/2Q^* mice over ten minute exploration in the open field (top) or during walking (black circles) and non-walking states (grey circles) (bottom). N = 5 wild-type and N = 6 *Atxn1^154Q/2Q^* mice One-way ANOVA with Tukey’s post hoc * p< 0.05, *** p < 0.001.

To determine the effect in the context of a cerebellum-associated motor behavior, we examined how SCA1 impacts the PCs activity in these lobules during walking and non-walking states. In anterior lobule IV of SCA1 mice, we found a significant reduction in PC peak amplitude and AUC only during the non-walking state (Figure 9A, bottom). On the other hand, in posterior lobule VI, PC activity was altered both during walking and non-walking states (Figure 9B, bottom). During walking, the frequency and AUC of SCA1 PCs were increased compared to WT controls. At rest, however, the amplitude of the PC calcium events was decreased in lobule VI of SCA1 mice.

Finally, we examined the impact of disease on the regional differences in PC activity during walking. In contrast to regionally-distinct mechanism of increase in PC activity during walking state in wild-type mice, the anterior and posterior PCs of SCA1 mice used similar mechanism and increased frequency, amplitude and AUC during walking.

These results indicate intrinsic regional differences in wild-type PC activity between the anterior and posterior cerebellum during self-motivated unrestricted walking and non-walking states. Moreover, in SCA1 mice, we found profound changes in PC activity in posterior lobule VI implicating regional vulnerability at a functional level. Finally, SCA1 impacted regional differences in PC activity

## Discussion

The advancement of sequencing techniques, including single-cell and spatial transcriptomics has raised the awareness of the regional heterogeneity of gene expression in healthy brain. Our results underscore the importance of intracerebellar regional identity and provide insight into a complex interplay between regional heterogeneity and regional vulnerability in neurodegeneration.

We identified a surprising number of DEGs in a comparison of the anterior and posterior wild-type cerebellum, a result supportive of intracerebellar heterogeneity. We found that regional variability can be seen at the level of neuronal activity. Specifically, we found regional differences in PC activity during walking and non-walking states in wild-type mice. In both regions, PC activity was significantly increased during walking, perhaps reflective the role of cerebellum in the coordination of free movement. However, while the increased PC activity of the anterior lobule IV during walking was largely characterized by an increased frequency of calcium events, in lobule VI it was observed as an increase in the amplitude and AUC. An increased frequency of calcium events points to one mechanism of signaling, while an increase in amplitude/AUC points to another mechanism, supporting functional intracerebellar heterogeneity. Future studies could explore relationship between genes and pathways enriched in anterior and posterior regions and these regional differences in function, to identify molecular underpinnings of functional heterogeneity.

Notably, we have also found that cerebellar disease impacts regional heterogeneity both at the level of gene expression and neuronal activity. It is possible that the reduction in heterogeneity allows for disease pathogenesis to spread to other regions, creating a vicious cycle..

We identified a pattern of exacerbated pathology across the anterior to posterior axis of the cerebellar vermis of SCA1 mice, consistent with observations of selective PC vulnerability in the posterior vermis of patients with SCA1. We found that while many pathways were altered in both regions, the posterior cerebellum exhibited larger number of affected pathways. This result may indicate that the number of affected pathways, instead of number of DEG or a specific pathway, underlie increased vulnerability. This is consistent with many studies showing almost complete rescue of disease symptoms by manipulating different pathways.

Finally,

Increased vulnerability in disease is usually defined as most severe neuronal pathology in particular brain region. Some pathological changes such as neuronal atrophy can be compensatory mechanisms to preserve function in disease.An important question is whether there is increased vulnerability at the functional level. In SCA1 mice, we found that posterior PCs are more impacted at the level of function as well as at the level of pathology. In particular, our results show that the activity of posterior PCs was altered both during walking and non-walking states, while anterior PC were altered only during rest. Future studies will determine whether this is also the case in other brain regions and other diseases.

Intracerebellar differences in degeneration have been described in other cerebellar diseases. In Spinocerebellar Ataxia type 5 (SCA5) posterior lobules of the cerebellum to preferentially undergo degeneration (Perkins et al., 2016). However, the posterior cerebellum is not universally more vulnerable. In fact, the opposite pattern of cerebellar degeneration is seen in Niemann-Pick disease, type C1 (NPC1), where the anterior vermis is more affected (Martin et al., 2019). Thus, it is likely that the region-specific effect of a disease is not simply a question of one region being generally more vulnerable than another, but a factor of the disease-specific mechanism and the underlying molecular signature of that cerebellar region. The results of these studies and the work presented here highlight the importance of assessing future disease-associated molecular changes in a region-specific manner, as bulk RNAsequencing is likely to mask any differences across regions that might confer vulnerability.

While these results indicate an exacerbated change in the cerebellar transcriptome and activity in posterior lobules, they are limited to one stage of a progressively worsening disease. Our histological analyses show that the posterior cerebellum worsens with time and that the anterior cerebellum begins to show signs of declining PC health at later time points. Thus, it is likely that transcriptomic analysis would demonstrate evidence of PC dysfunction and pathology with aging. Future transcriptomic work following the progression of SCA1 in both regions over time would thus be useful for understanding both how the disease uniquely affects these two regions and what mechanisms may be useful for delaying cerebellar dysfunction in vulnerable regions.

Our findings advance understanding of cerebellar physiology and neurodegeneration in several ways. We characterized cerebellar heterogeneity at the level of the gene expression and PC activity during unrestrained walking in the wild-type mice. These results increase our understanding of cerebellar physiology and provide a foundation on which to test genes and pathways that could contribute to the observed differences in PC activity and underlie increased regional-vulnerability in cerebellar disease. Our results show that intraregional analysis may be needed to identify differences in disease susceptibility and underlying molecular processes that may mitigate disease. This work provides targets for further exploration of cerebellar heterogeneity and the impact its loss may have on cerebellar disease. We hope that the database of genes and pathways altered in a regional manner will inspire studies of increased disease vulnerability or resistance to provide targets for successful treatments for SCA1 and other cerebellar diseases.

## Figure legends

**Supplementary Figure 1. Molecular layer thickness decrease is not uniform across cerebellar vermis and hemisphere in 18-week-old *Atxn1^154Q/2Q^* mice A.** Sections from 18 weeks old *Atxn1^154Q/2Q^* mice and wild-type controls were stained for calbindin, marker of Purkinje cells. Wide field images were acquired using an epifluorescent microscope. WT images (top), *Atxn1^154Q/2Q^* (bottom), vermis sections (left), hemisphere sections (right). **B.** Percent loss of molecular layer thickness compared to wildtype was calculated per lobule and combined into cerebellar cortical regions. Anterior vermis (top, lobules I/II-IV/V), posterior vermis (middle, lobules VI-X), and hemispheres (bottom) n = 4 per genotype, WT in black bars, *Atxn1^154Q/2Q^* in pink bars, data are presented as mean ± SEM, Unpaired t-test, ** p<0.05, **** p<0.0001.

**Supplementary Figure 2. Assessment of dendritic atrophy, synaptic loss, and gliosis in four vermal regions in 12-week-old *Atxn1^154Q/2Q^* mice. A.** Representative Calbindin IHC images of lobule II (top) and X (bottom) for WT (left) and *Atxn1^154Q/2Q^* (right); used to measure molecular layer width and calbindin intensity. **B.** Measurement of molecular layer thickness. n = 3-4 WT (black bars), 4-5 *Atxn1^154Q/2Q^* (pink bars), unpaired t-test, * p < 0.05. **C.** Representative Calbindin and VGLUT2 IHC images of lobule II and X, WT (top) and *Atxn1^154Q/2Q^* (bottom), Calbindin staining (left), VGLUT2 staining (middle), merged images (right). Images used to take ratio measurements between molecular layer width and VGLUT2 puncta width. Example measurements drawn in magenta and green lines on the merged images. **D.** Calbindin staining intensity measurements in the four vermal lobule regions. n = 3-4 WT (black bars), 4-5 *Atxn1^154Q/2Q^* (pink bars), unpaired t-test, * p < 0.05. **E.** Ratio of VGLUT2 puncta/ molecular layer width measurements in the four vermal lobule regions. n = 3-4 WT (black bars), 4-5 *Atxn1^154Q/2Q^* (pink bars), unpaired t-test, * p < 0.05, ** p < 0.005. **F.** Representative GFAP IHC images of lobule II (top) and X (bottom) for WT (left) and *Atxn1^154Q/2Q^* (right); used to measure intensity of GFAP staining normalized to WT images. **G.** GFAP staining intensity measurements in the four vermal lobule regions. n = 4-5 WT (black bars), 6-7 *Atxn1^154Q/2Q^* (pink bars), unpaired t-test, * p < 0.05. **H.** Representative Iba1 IHC images of lobule II (top) and X (bottom) for WT (left) and *Atxn1^154Q/2Q^* (right); used to measure density of microglia in the molecular layer. I. Iba1 density measurements in the molecular layer of four vermal lobule regions. n = 3 WT (black bars), 4 *Atxn1^154Q/2Q^* (pink bars), unpaired t-test. Data are presented as mean ± SEM.

**Supplementary Figure 3. Cerebellar dissection and RNAsequencing analysis method. A.** Cartoon of cerebellar dissection. Cerebellar vermis sections are dissected into anterior (CCAV) and posterior regions (CCPV) at primary fissure (red line). Deep cerebellar nuclei were punched out (red circles) to isolate the cerebellar cortex of hemispheres (CCH). Tissue from each region is collected and saved separately. RNA is extracted from each region separately to be used for RNAsequencing and RTqPCR. **B.** Schematic of RNAsequencing analysis method. Comparisons between WT and *Atxn1^154Q/2Q^* mice in each cerebellar cortical region (red arrows) producing differentially expressed genes (DEGs) in *Atxn1^154Q/2Q^* mice. Comparisons within WT cerebellar cortical regions (yellow arrows) producing DEGs between each cerebellar cortical region in wild-type mice. Comparisons within *Atxn1^154Q/2Q^* cerebellar cortical regions (pink arrows) producing DEGs between each cerebellar cortical region in the *Atxn1^154Q/2Q^* mice.

**Supplementary Figure 4. GO pathway analysis of differentially expressed genes across cerebellar cortical regions in *Atxn1^154Q/2Q^* mice. A.** Enriched pathways based on GO: Biological process. **B.** Enriched pathways based on GO: Molecular Function. For each analysis, the top 500 DEGs based on absolute value of logFC between WT and *Atxn1^154Q/2Q^* mice in each cerebellar cortical region were run through the GO database. Most pathways were enriched in all cerebellar cortical regions. Dot size represents number of genes in the enriched pathway, and color of the dot represents p-value.

**Supplementary Figure 5. Heterogeneity in SCA1 mice. A-C.** Heatmaps of top up and down regulated DEGs in between the anterior and posterior vermis (A), the anterior vermis and the cerebellar hemispheres (B), and the posterior vermis and cerebellar hemispheres (C). Heatmaps were generated by ranking the top 15 up and down regulated DEGs based on logFC value and plotting the counts per million (CPM) of each gene in individual samples. Expression is normalized per gene in the heatmap representation. D. KEGG pathway analysis of enriched pathways of top 500 DEGs between the anterior and posterior vermis, the anterior vermis and the cerebellar hemispheres, and the posterior vermis and cerebellar hemispheres. Dot size represents number of genes in the enriched pathway, and color of the dot represents p-value.

