## Supplementary figures and images for "Loss of intracerebellar heterogeneity and selective vulnerability in Spinocerebellar ataxia type 1 neurodegeneration"

### Supplementary Figures

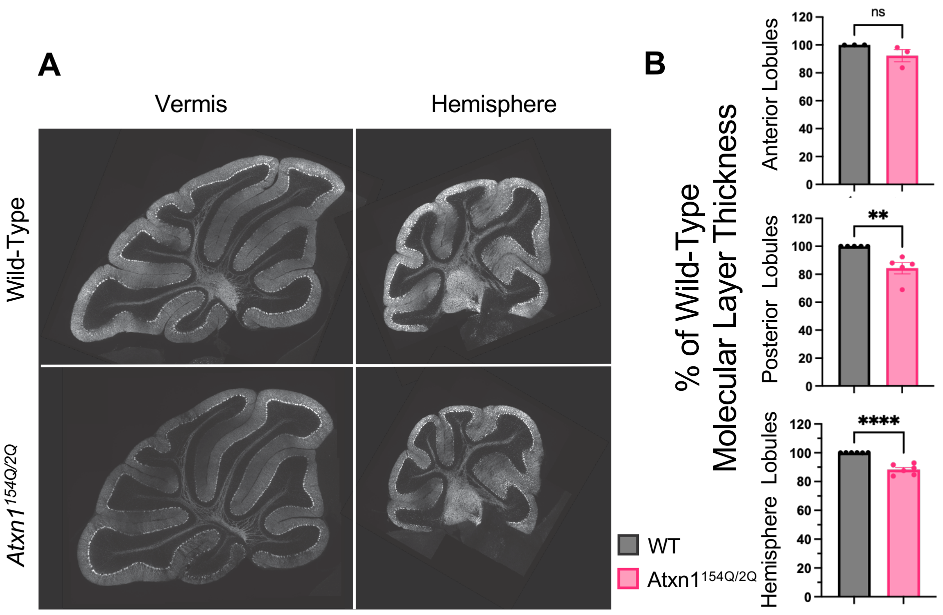

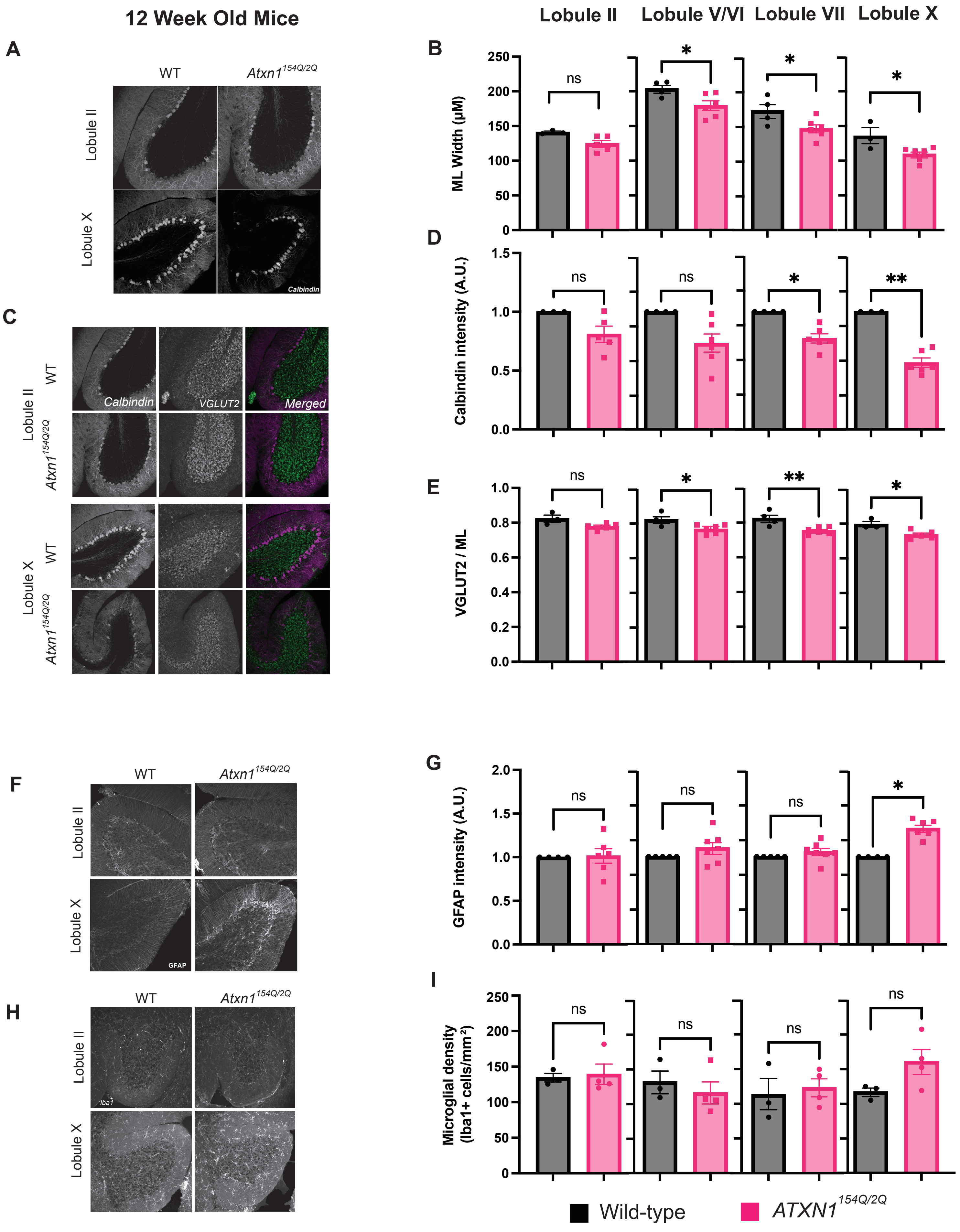

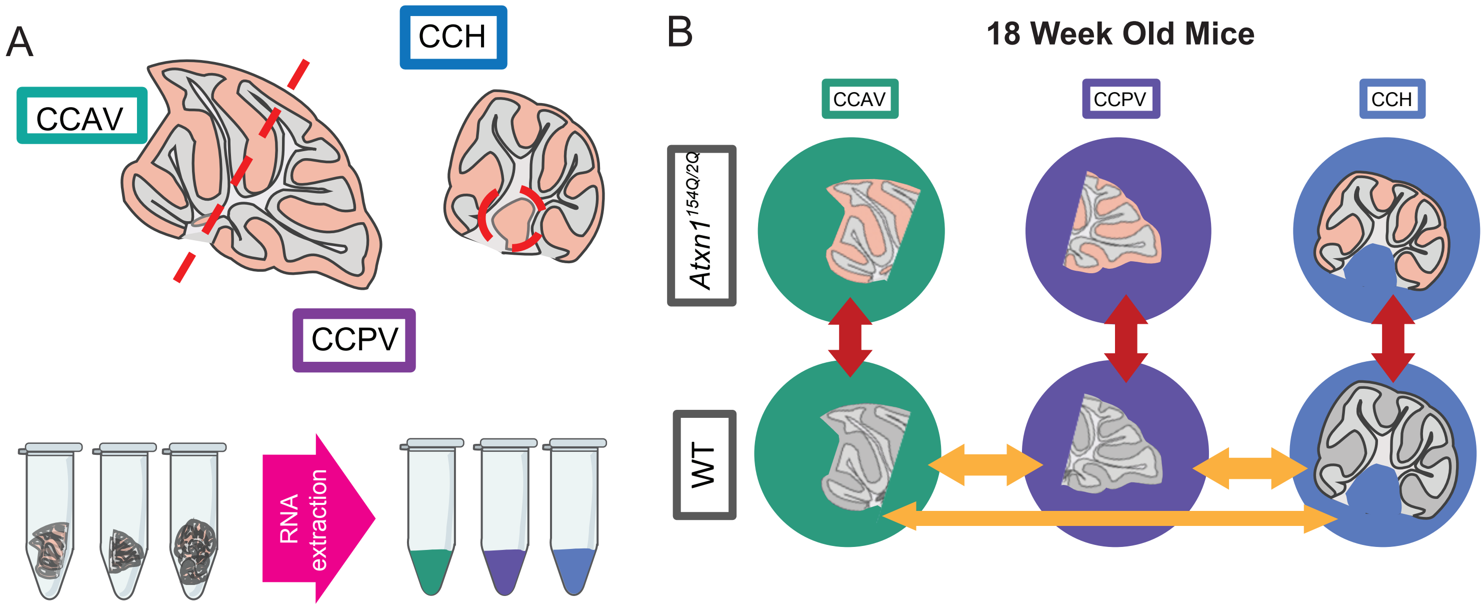

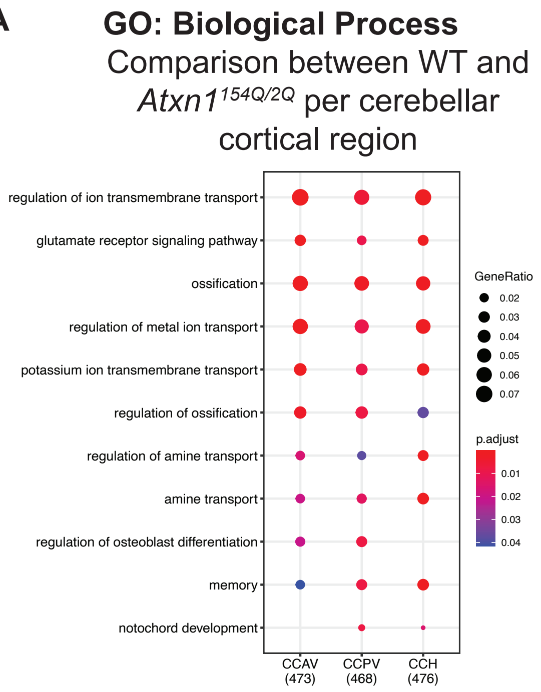

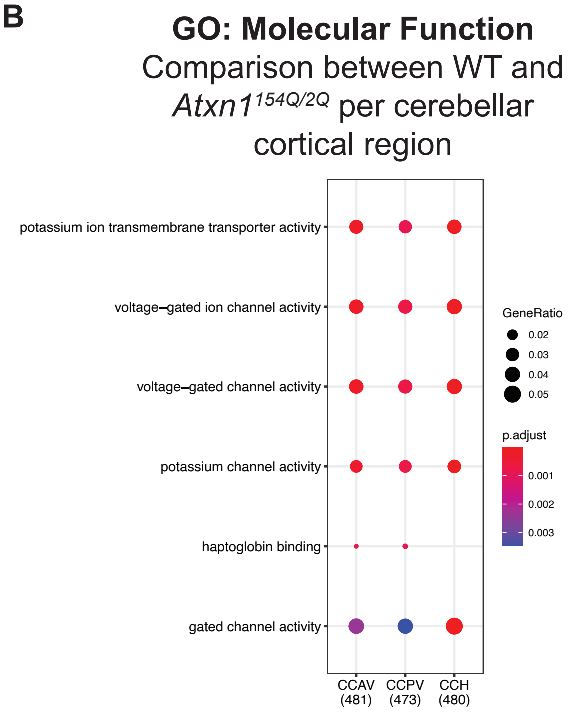

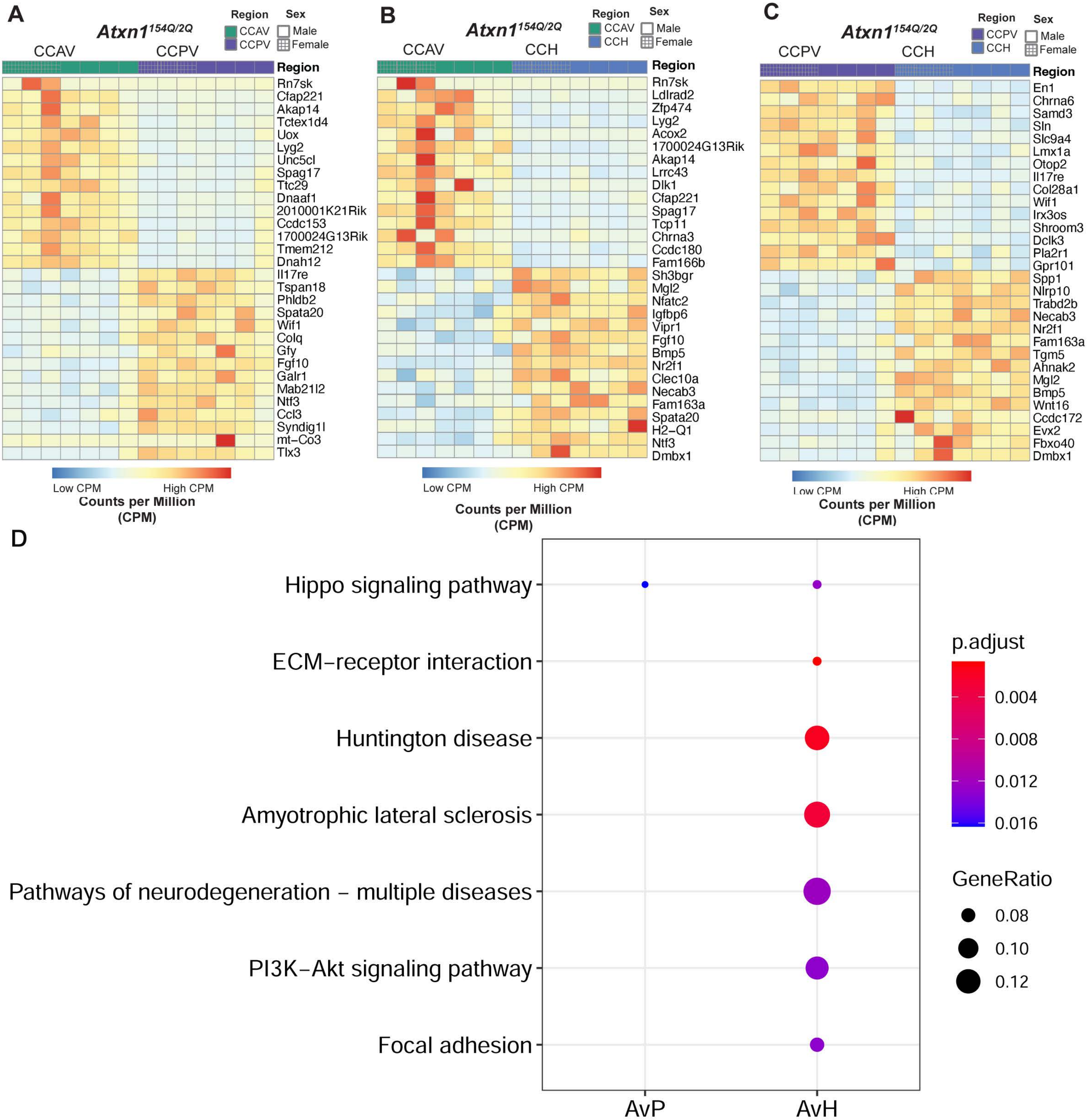
